# Geographically structured genomic diversity of non-human primate-infecting *Treponema pallidum* subsp. *pertenue*

**DOI:** 10.1101/848382

**Authors:** Benjamin Mubemba, Jan F. Gogarten, Verena J. Schuenemann, Ariane Düx, Alexander Lang, Kathrin Nowak, Kamilla Pléh, Ella Reiter, Markus Ulrich, Anthony Agbor, Gregory Brazzola, Tobias Deschner, Paula Dieguez, Anne-Céline Granjon, Sorrel Jones, Jessica Junker, Erin Wessling, Mimi Arandjelovic, Hjalmar Kuehl, Roman M. Wittig, Fabian H. Leendertz, Sébastien Calvignac-Spencer

## Abstract

**Background:** Increasing evidence suggests many non-human primate (NHP) species in sub-Saharan Africa are infected with *Treponema pallidum* subsp. *pertenue* (TPE), the bacterium causing yaws in humans. In humans, yaws is characterized by lesions of the extremities and face, while *Treponema pallidum* subsp. *pallidum* (TPA) causes venereal syphilis and is characterized by primary lesions on the genital, anal or oral mucosae, and has not been detected in NHPs. Due to a paucity of genetic data, it remains unclear whether other *Treponema pallidum* (TP) subspecies found in humans also occur in NHP and how the genomic diversity of TPE lineages that do occur in NHPs is distributed across hosts and space.

**Methodology:** We observed a combination of yaws- and syphilis-like symptoms in sooty mangabeys (*Cercocebus atys atys*) in Taï National Park (TNP), Côte d’Ivoire and collected swabs and biopsies from symptomatic animals. We also collected NHP bones from eight species from TNP, as well as from 19 species at 12 field sites across sub-Saharan Africa. Samples were screened for TP DNA using PCRs. In-solution hybridization capture coupled with high throughput sequencing was used to sequence TP genomes from positive samples.

**Principal findings:** We generated four nearly complete TP genomes from biopsies and swabs and five partial genomes from bones. Phylogenomic analyses revealed that both syphilis- and yaws-like lesions of sooty mangabeys within a single social group in TNP were caused by TPE. All TPE genomes determined from these sooty mangabeys were different and exhibited divergence levels not observed in TPE from a single species at any other field site where the disease seems to be epizootic. In general, simian TPE isolates did not form monophyletic clades based on host species or the type of symptoms caused by an isolate, but rather clustered based on geography.

**Conclusions:** There is a large diversity of TPE strains infecting NHPs in TNP. Our observations within a single social group of sooty mangabeys do not support the epizootic spread of a single clone, but rather points towards frequent independent introductions of the bacterium, which can cause syphilis- and yaws-like lesions. On a larger scale, the geographic clustering of TPE genomes might be compatible with cross-species transmission of TPE within ecosystems or environmental exposure leading to acquisition of closely related strains.

**Author’s summary:** Individuals in several populations of wild non-human primates (NHP) in sub-Saharan Africa are infected with *Treponema pallidum* subsp. *pertenue* (TPE), a pathogen causing yaws disease in humans. In humans, yaws is characterized by skin lesions of the extremities and face. In contrast, *Treponema pallidum* subsp. *pallidum,* which causes venereal syphilis in humans, has not been observed in NHPs. We describe a combination of yaws- and syphilis-like symptoms in a sooty mangabey (*Cercocebus atys atys*) social group in Taï National Park (TNP), Côte d’Ivoire. We sampled lesioned animals and collected and tested NHP bones from field sites across sub-Saharan Africa. We were able to reconstruct four genomes from swabs/biopsies and five partial genomes from bone samples. Phylogenomic analyses revealed that syphilis-like lesions and yaws-like lesions in TNP were caused by a large diversity of TPE strains. Additionally, simian TPE isolates did not form monophyletic clades based on the host species or the types of symptoms caused by an isolate, but rather clustered by geographic origin. This is suggestive of cross-species transmission of TPE within ecosystems or environmental exposure leading to acquisition of closely related strains.

## Introduction

Spirochete bacteria belonging to the species *Treponema pallidum* (TP) have affected mankind since at least the late 15^th^ century [1] and cause a large global disease burden in humans [2, 3]. Three pathogenic subspecies are currently recognised [4] that are morphologically similar, but distinguishable genetically, epidemiologically, and clinically into three disease syndromes; yaws (subsp. *pertenue;* TPE), venereal syphilis (subsp. *pallidum;* TPA) and bejel (subsp. *endemicum;* TEN) [5, 6]. Though treatable, these treponematoses remain major public health threats across the globe [2–4]. Annually, nearly 8 million new cases of venereal syphilis are reported globally [7], while in the 13 countries where yaws remains endemic, it is estimated that more than 80,000 new cases occur. Exact estimates of the number of bejel cases from the Sahel region and Arabian Peninsula are not available [8, 9]. Efforts to reduce the prevalence of these trepanomatoses are underway, particularly for yaws, for which an ongoing campaign aims to eradicate the disease globally by 2030 [10].

Key questions for the potential success of eradication efforts and for understanding TP evolution is the degree to which other animals are infected with these pathogens and whether cross-species transmission occurs. To date, only TPE has been shown to infect NHPs [11–13]. The TPE strains infecting humans and NHPs are extremely similar, and there is no clear evidence for phylogenetic separation of NHP-infecting and human-infecting TPE strains, as strains do not form well-supported reciprocally monophyletic groups [12, 13]. Whether between species transmission events have occurred between different NHP species or humans remains unclear due to a paucity of genomic data from both humans and wildlife. However, in an experimental setting, the simian Fribourg-Blanc strain isolated from Guinea baboons (*Papio papio*) induced classical yaws symptoms in humans [14]. Similarly, human infecting TP strains were reported to elicit yaws-like symptoms in NHPs [15]. This suggests that molecular compatibility barriers to between-species transmission of TPE are low, though other barriers to spillover might exist [16].

Cross-species transmission events are governed by a number of factors that include; spatial proximity of the hosts in an ecosystem, phylogenetic relatedness of the hosts, and the biology of both the host and the pathogen [17]. One tool for assessing the potential of between-species transmission for a pathogen is to look for signals of historic transmission in the phylogeny of a pathogen [18]. This often takes the form of testing for incongruence between the host phylogeny and that of a pathogen [19, 20]. Such analyses have not been formally attempted for TPE, in large part due to a lack of genomic data from TPE infecting both humans and wildlife. Data on the genetic diversity of TPE in a diversity of NHP hosts could reveal historic between-species transmission events and provide data that would enable the detection of zoonotic transmission events between NHPs, and between NHP and humans, if any at all are occurring.

Here, we describe a combination of yaws- and syphilis-like symptoms in a single group of sooty mangabeys (*Cercocebus atys atys*) in Taï National Park (TNP), Côte d’Ivoire and test whether these diverse symptoms were caused by a single lineage of a TP subspecies. In addition, to examine whether cross-species transmission of TP occurs between NHPs in a single ecosystem, we drew on recent findings that TP can be detected in NHP bones [21] to screen NHP bones from TNP. To enable us to further test whether the phylogenies of TP subspecies mirror that of their hosts, we also analysed non-symptomatic bones from twelve species sampled across eight sub-Saharan countries (Table S2).

## Materials and Methods

### Ethical statement

All procedures performed on the sooty mangabeys in TNP were approved by the Ministry of Environment and Forests as well as the Ministry of Research, the Office Ivoirien des Parcs et Réserves, and the director of TNP. A veterinarian carried out all sampling following good veterinarian practice. Animal welfare was considered in all procedures carried out and anesthetized animals were monitored for vital functions and remained under close supervision from the time of induction until full recovery and until the animals were able to reunite with their social group.

### Study sites and samples

In January 2014, sooty mangabeys from a habituated social group in TNP were observed with orofacial lesions and lesions of their distal extremities; two animals had been previously sampled and TPE was determined to be the cause of the infection [12]. Over the next two years, other individuals started showing genital ulcerations and necrotizing dermatitis on the inner parts of the thighs and ventral abdomen, often with visible yellow crusts. Orofacial and genital lesions were still observed on other animals in the group during the study period (Figure 1). Three more individuals with visible lesions were chemically immobilised using a combination of xylazine (1 mg/kg) and ketamine (10 mg/kg) administered by blowpipe. Biopsy and lesion swab samples were collected from genital and orofacial lesions (Table 1). Samples for molecular biology analysis were preserved in RNAlater® (Life technologies, NY) and shipped to the Robert Koch Institute, Berlin, Germany for laboratory analysis.

**Figure 1:**
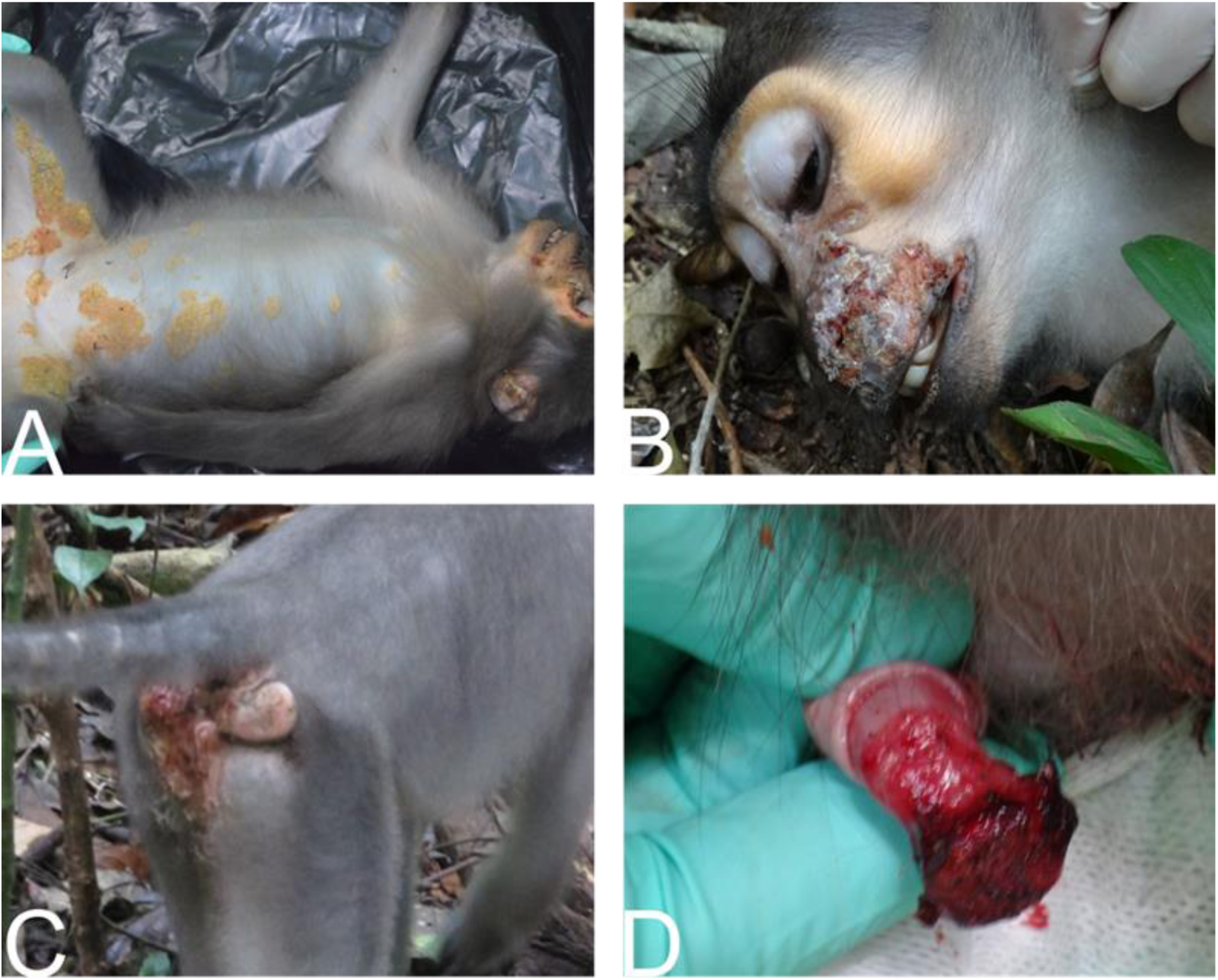
Lesions due to TPE infection in sooty mangabeys. **A.** Necrotizing dermatitis of inner parts of the thighs and ventral abdomen with yellowish crusts **B.** Necrotic orofacial lesions **C.** genital lesions in females **D.** Genital lesions in males

**Table 1:**
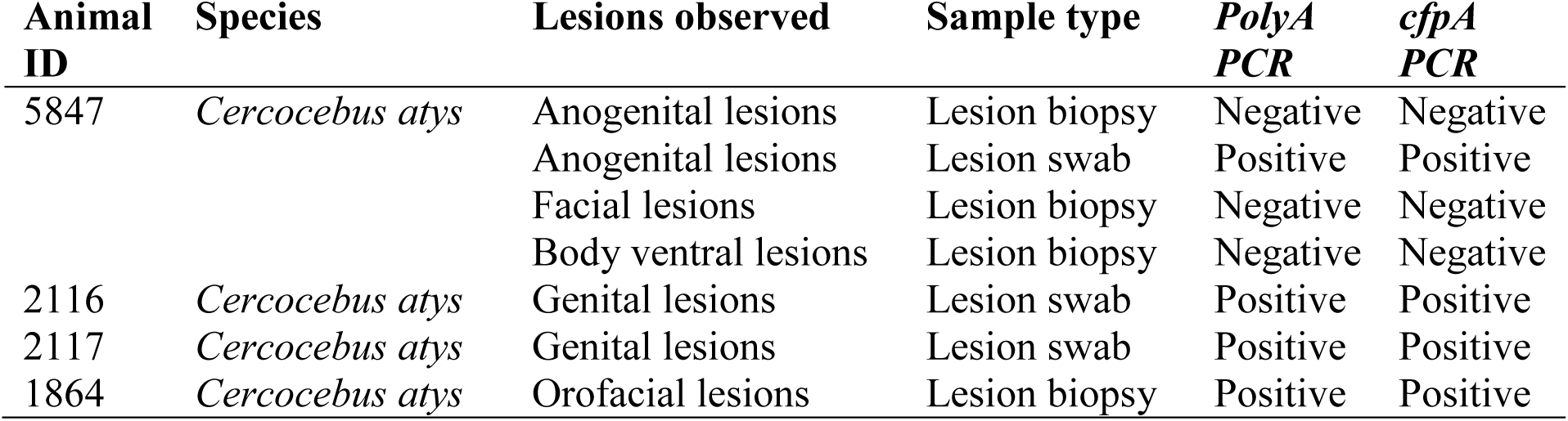
Lesion and sample types observed at TNP and PCR screening results for each sample.

To get insights into the NHP trepanomatoses circulating in TNP during the last three decades, we screened non-symptomatic bones collected opportunistically at TNP (*N*=67; Table S2). We also screened NHP bones collected from 11 additional field sites in sub-Saharan Africa (*N*=80; Table S2) collected through the support of the Pan African Program: The Cultured Chimpanzee (PanAf; www.panafrican.eva.mpg.de). NHP bones were assigned to particular species using molecular methods targeting the 16S gene as described previously [22], coupled with morphological assignment by experts in the field.

### DNA extraction

DNA was extracted from both skin biopsies and swabs using the DNeasy Blood and Tissue kit (QIAGEN, Germany) following the manufacturers’ instructions. DNA was extracted from bones using a silica-based method. Briefly, non-lesioned bones were drilled using a fine drill bit at slow speed to produce ∼150 mg of bone powder. The drilling was done in a designated sealed glove box, both to prevent any contamination of the bones and to prevent exposure of researchers to pathogens that might be present in the bones (e.g., in TNP we have many cases of sylvatic anthrax (*Bacillus cereus* biovar *anthracis*), which can be cultured from bones) [23]. The box was UV sterilized and subsequently surfaces were bleached following the drilling of each bone and extraction. Drill bits were changed on every bone to prevent cross contamination. The DNA extraction from the bone powder was done according to the protocol described in detail in [24] following a modified protocol of Rohland and Hofreiter, 2007 and Gamba *et al.* 2015 [25, 26]. Extracted DNA from all experiments was quantified with a Qubit fluorometer with the double stranded DNA high sensitivity kit (Invitrogen™, Germany) following the manufacturer’s instructions. DNA was subsequently stored at −20°C.

### Screening for Treponema pallidum

To screen for *Treponema pallidum* DNA in swabs and biopsy samples, we performed an end-point polymerase chain reaction (PCR) assay targeting the 67 bp of the *polyA* gene fragment, using primers (Table S1) previously developed for screening human clinical specimens [27]. PCR reactions were performed in 25µL reactions; up to 200 ng of DNA was amplified using 1.25 U of high-fidelity Platinum Taq™ polymerase (Invitrogen®, Germany), 10x PCR buffer (Invitrogen®, Germany), 200µM dNTPs, 4mM MgCl2, 200nM of both forward and reverse primers. The thermal cycling profile was as follows; denaturation at 95°C for 5 min, followed by 40 cycles of 95°C for 15 sec, 60°C for 30 sec, and 72°C for 1 min. An elongation step at 72°C for 10 min was included. Positive and negative controls were included.

The 67 bp amplified product is too short for direct Sanger’s sequencing, so to confirm the results of this initial screening, positive samples were further tested with a semi-nested assay targeting the cytoplasmic filament protein gene (*cfpA*) [28]. These primer pairs (Table S1) yielded a 352 bp outer product in the primary PCR and a 189 bp inner fragment in the nested PCR. In both primary and nested PCR assays, the reactions were performed as follows; about 200 ng of DNA was amplified in a 25µL reaction using 1.25 U of high-fidelity Platinum Taq™ polymerase® (Invitrogen®, Germany), 10x PCR buffer (Invitrogen®, Germany), 200µM dNTPs, 4mM MgCl2, 200nM of both forward and reverse primers. Subsequently, 2 µl of a 1:20 dilution of the first round PCR product was used as input template for the nested PCR. The thermal cycling profile for both the primary and nested PCRs were the same as in the initial screening assay.

PCR products were visualized on a 1.5% agarose gel stained with GelRed® (ThermoFisher scientific, Germany). Positive bands were purified using the PureLink™ Quick Gel Extraction Kit (Invitrogen®, Germany) following the manufacturer’s protocol. Purified products were stored at −20°C until they were sequenced using using BigDye Terminator v3.1 Cycle Sequencing Kit and sequences were compared to publicly available sequences in EMBL through BLAST [29]. All samples that tested positive from the confirmatory assay were selected for whole genome in-solution hybridization capture and high throughput sequencing.

Potential DNA degradation in bone samples precluded the use of the above mentioned confirmation assay. To select the most promising samples, we estimated copy numbers in the bones, using a real-time qPCR also targeting the 67bp fragment of the *polA* gene [27]. Samples were tested in duplicate. Briefly, 5 µl of total DNA was amplified in a 25µL qPCR reaction containing 10x PCR buffer (Invitrogen®, Germany), 200µM dNTPs, 4mM MgCl2, 300nM of both forward and reverse primers, 100nM of the probe and 0.5U of high-fidelity Platinum Taq™ polymerase® (Invitrogen®, Germany). The thermal cycling profile was set as follows; DNA denaturation at 95°C for 10 min followed by 45 cycles at 95°C for 15s and 60°C for 34s. All bone samples that had detectable TP DNA in duplicate reactions were selected for in-solution hybridization capture and high throughput sequencing.

### Library preparation, genome wide capture and high throughput sequencing

For tissue and swab sample extracts, we sheared 1000 ng of DNA per sample to 400 bp fragments using the Covaris S2 (intensity: 4, duty cycle: 10%, cycles per burst: 200, treatment time: 55 seconds and temperature: 4-5°C). Bone samples were not sheared due to the potentially fragmented nature of DNA in these older specimens. Following shearing, two library preparation methods were followed. In one method, the DNA was then converted into double indexed Illumina libraries using the NEBNext® Ultra™ II DNA Library Prep kit (New England Biolabs®; samples: Tai_105, Boe_092, 2117, 5847, 1864). In the other, the Accel-NGS (Swift BIOSCIENCES™; samples: 11786, 15028, 22_52, 2116) was used to convert the DNA into single indexed Illumina libraries following the respective manufacturer’s protocols (Table 2). Samples 11786, 15028 and 1864 were sequenced previously [12, 21], but in this study, we generated new libraries for sample 1864 to improve genome coverage for this sample. All generated libraries were quantified using the KAPA library quantification kit (KAPA Biosystems, SA) following the manufacturer’s instructions. In all library preparations, chicken DNA was included as a control.

**Table 2:** Summary table for library preparation, capture and sequencing methods used for each sample.

| Sample ID | NHPs species | Sample type | Library Prep method |  | Capture method |  | Sequencing method | NP / Country |
| --- | --- | --- | --- | --- | --- | --- | --- | --- |
|  |  |  | NEBNext | Accel NGS | In-solution | Microarray |  |  |
| **Boe_092** | *Cercocebus atys* | bone | X |  | X |  | Miseq | Boe/GB |
| **2117** | *Cercocebus atys* | swab-genital lesion | X | X | X |  | Miseq | TNP/CI |
| **5847** | *Cercocebus atys* | swab-genital lesion | X |  | X |  | Miseq | TNP/CI |
| **2116** | *Cercocebus atys* | swab-genital lesion |  | X | X |  | Miseq | TNP/CI |
| **1864** | *Cercocebus atys* | biopsy-face lesion | X | X | X |  | Miseq | TNP/CI |
| **control** | chicken DNA | tissue | X |  | X |  | Miseq | - |
| **11786** | *Pan troglodytes verus* | bone |  | X |  | X |  | TNP/CI |
| **15028** | *Pan troglodytes verus* | bone |  | X |  | X |  | TNP/CI |
| **Tai_105** | *Piliocolobus badius* | bone | X |  | X |  | Miseq | TNP/CI |
| **22_52** | *Piliocolobus badius* | bone |  | X | X |  | Miseq | TNP/CI |

NP: National Park, GB: Guinea Bissau, TNP: Tai National Park, CI: Côte d'Ivoire

Libraries were enriched for *Treponema pallidum* sequences using in-solution hybridization capture with biotinylated RNA baits and following the manufacturer’s protocol (myBaits, Arbor Biosciences). The baits spanned the simian derived Fribourg-Blanc reference genome (RefSeq ID: NC_021179.1) with a 2x tiling. In-solution hybridization capture was done for two rounds of 48 hrs each. After each round of capture, a post capture amplification step was performed using the KAPA HiFi master mix (KAPA Biosystems, SA) with P5 and P7 Illumina primers to generate about 200 ng of enriched DNA per sample. The post-capture amplification thermal profile was as follows: initial hot start at 98°C for 2 min followed by 12 cycles or what is required to generate 200 ng at 98°C for 20 sec, 65°C for 30 sec and 72°C for 45 sec. Enriched libraries were quantified using the KAPA library quantification kit (KAPA Biosystems, SA). Prior to sequencing, libraries were diluted to 4 nM and pooled for sequencing on an Illumina Miseq with 300 bp paired end reads (V3 chemistry; Table 2) at the Robert Koch Institute (Berlin, Germany).

### Bioinformatics analysis

Paired-end reads generated here, along with available SRAs from prior TP sequencing efforts (Table 3 and Table S3) were trimmed using Trimmomatic v0.38, removing the leading and trailing reads below Q30, clipping any part of the read where the average base quality across 4 bp was less than 30, and removing surviving reads less than 30 bp in length [30]. Surviving read pairs were merged using Clip and Merge version 1.7.8 with default settings [31]. Merged reads and surviving singletons were combined and mapped to TPE Fribourg-Blanc (RefSeq ID: NC_021179.1) using BWA-MEM [48] with a minimum seed length of 29. Mapped reads were sorted using Picard’s SortSam and subsequently de-duplicated with Picard’s MarkDuplicates (https://broadinstitute.github.io/picard/index.html). Alignments with a MAPQ smaller than 30 and a mapping length lower than 30 were also removed using SAMtools [32]. Consensus sequences were then called in Geneious v11.1.5 [33], and theimportance of the minimum coverage requirement and identity threshold was explored by requiring a minimum coverage of 3, 5, and 10 unique reads (i.e., 3X, 5X or 10X), and a threshold of 50% or 95% identity for a base to be called, resulting in a total of six different consensus sequences called for each sample.

**Table 3:** Mapping results for non-human primate *Treponema pallidum* subsp. *pertenue* strains from TNP and BNP. Shown is the genome coverage at both 1X and 3X.

| Sample ID | NHPs species | Sample type | Deduplicated reads mapped | RefSeq length | Pos. covered 1X | Pos. covered 3X | % genome 1X | % genome 3X |
| --- | --- | --- | --- | --- | --- | --- | --- | --- |
| **1864** | *Cercocebus atys* | biopsy-face lesion | 202416 | 1140481 | 1116210 | 1109692 | 97.87 | 97.30 |
| **Boe_092** | *Cercocebus atys* | bone | 36407 | 1140481 | 793392 | 440040 | 69.57 | 38.58 |
| **2117** | *Cercocebus atys* | swab-genital lesion | 44143 | 1140481 | 1009366 | 778675 | 88.50 | 68.27 |
| **5847** | *Cercocebus atys* | swab-genital lesion | 457063 | 1140481 | 1117566 | 1115883 | 97.99 | 97.84 |
| **2116** | *Cercocebus atys* | swab-genital lesion | 131550 | 1140481 | 1119023 | 1102573 | 98.12 | 96.68 |
| **Control** | chicken | tissue | 1 | 1140481 | 52 | 0 | 0.005 | 0.00 |
| **11786** | *Pan troglodytes verus* | bone | 550 | 1140481 | 3362 | 525 | 0.29 | 0.05 |
| **15028** | *Pan troglodytes verus* | bone | 4409 | 1140481 | 81049 | 33410 | 7.11 | 2.93 |
| **Tai_105** | *Piliocolobus badius* | bone | 42719 | 1140481 | 867318 | 655599 | 76.05 | 57.48 |
| **22_52** | *Piliocolobus badius* | bone | 43373 | 1140481 | 711608 | 503757 | 62.40 | 44.17 |

Whole genome alignments were performed on each set of consensus sequences using the multiple sequence alignment program MAFFT [34]. In addition, we removed all putative recombinant genes [35] and selected conserved blocks using the Gblocks tool [36] in SeaView v4[37]. Phylogenetic inference was performed on the resulting alignments of informative positions after stripping off of all ambiguities and identical sites in the final data sets: 3X, 50%; (2726 positions); 3X, 95% (2726 positions); 5X, 50% (3383 positions); 5X, 95% (2726 positions); 10X, 50% (3203 positions) and 10X, 95% (2686 positions).

Final alignments were then uploaded to the online ATGC PhyML-SMS tool (http://www.atgc-montpellier.fr/phyml-sms/) for construction of a maximum likelihood phylogeny using smart model selection [38] with Bayesian Information Criterion and subtree pruning and regrafting (SPR) for the tree improvement and otherwise using default settings. Branch robustness was estimated using Shimodaira-Hasegawa approximate likelihood ratio test (SH-like aLRT [39]). The maximum likelihood (ML) tree was then rooted using TempEst (version 1.5.1), which estimated the best-fitting root of these phylogenies using the heuristic residual mean squared function, which minimizes the variance of root-to-tip distances [40]. Evolutionary pairwise distance between TP strains was estimated from the ML phylogeny using the Patristic program [41].

To explore the robustness of our phylogenetic analysis, we also ran a Bayesian Monte Carlo Markov Chain (BMCMC) analyses with BEAST (version 1.10.4) using a strict clock model and a coalescent process assuming constant population size. We examined the output of multiple runs for convergence and appropriate sampling of the posterior using Tracer (version 1.7.1) [42] before merging runs using Log Combiner (version 1.10.4) [43]. The best representative tree was then identified from the posterior set of trees and annotated with Tree Annotator (version 1.10.4: distributed with BEAST). The resultant maximum likelihood and maximum clade credibility (MCC) tree files were edited using iTOL (https://itol.embl.de/) [44].

Due to insufficient genome coverage for several bone samples (11786, 15028, 22_52), which could not be reliably processed with our phylogenetic pipeline, we performed phylogenetic read placement using the evolutionary placement algorithm tool EPA-ng to determine the position of individual reads on the TP phylogeny [45]. Briefly, we selected filtered merged and singleton reads that mapped to TPE that were 80 bp or longer. Surviving reads were then aligned to the genomes used to build the MCC tree, using the Parsimony-based phylogeny-aware read alignment program (PaPaRa) [46]. The resulting PaPaRa alignment was then split into the query reads and the original alignment using the split function available in the EPA-ng toolkit [46]. To estimate the best fitting evolutionary model for the phylogenetic placement of the query reads, both the reference tree and reference alignment were evaluated using the RaxML-ng toolkit [47]. The best fitting model was then used to place query reads on the reference tree with the EPA-ng tool. The resulting EPA-ng jplace trees were visualised as heat-trees depicting the percentage of reads placed on each branch using the gappa toolkit [48]. The settings for the visualisation were as follows: every query read was treated as a point mass concentrated on the highest-weight placement and multiplicity of each query read set to 1.

## Results

### Screen PCR and whole genome capture

All symptomatic animals sampled with biopsies or swabs tested positive for *Treponema pallidum* in at least one of the sample types collected (Table 1). Sequences generated for the respective assays were all identical and a representative sequence was been uploaded to Zenodo (doi.org/10.5281/zenodo.3540499). In addition, based on the *polyA* gene qPCR assay, we detected TP DNA in NHP bones in the following field sites and species/subspecies: TNP in Côte d’Ivoire (*Cercopithecus diana, Pan troglodytes verus* and *Piliocolobus badius*), Bili-Uere in the Democratic Republic of Congo (*Pan troglodytes schweinfurthii),* Boe in Guinea Bissau (*Cercocebus atys*), Budongo in Uganda (*Cercopithecus mitis, Colobus guereza*), and in East Nimba and through the Nationwide survey in Liberia (*Pan troglodytes verus*; Table S2).

Including prior efforts [12], from the sooty mangabey study group in TNP, we were able to sequence four TPE genomes from biopsy and swab samples with 3X coverage of 88.5% - 98.3% of the genomes (Table 3). For sample 1864, which was collected from a sooty mangabey with facial lesions in TNP and was previously sequenced [12], we improved the 3X genome coverage from 82.39% to 97.3%. Further, we recovered partial TPE genomes from two *Pan troglodytes verus* (11786 and 15028) and two *Piliocolobus badius* (22_52 and Tai_105) bones from TNP, Côte d’Ivoire, as well as one *Cercocebus atys* bone from Boe in Guinea Bissau (range for all bone samples = 0.2 – 76% 1X genome coverage and 0.1 – 57% 3X genome coverage). The control library (Chicken DNA) had only a single read surviving the read quality control processing, translating to 0.005% 1X genome coverage.

### Phylogenetic analyses

Phylogenetic analysis of reconstructed genomes from this study and all other TPE and TEN genomes and a selection of TPA genomes in Genbank (Table S3) yielded tree topologies largely consistent in the PhyML-SMS and BEAST based approaches. The exception was the position of the TEN strains in the ML trees, which clustered with the TPE strains (Supplementary trees). To understand this contradictory finding, we performed an in-group analysis focussed on TPE and TEN strains. The ML trees were then rooted with TempEst (version 1.5.1) [40], which revealed that the root was located on the branch separating TPE and TEN, as observed previously [12] (supplementary trees).

Overall, the ML and MCC tree topologies resolved into distinct reciprocally monophyletic groups representative of the TP subspecies (TPA, TPE, and TEN). The TPE clade included both human and all NHP infecting strains, while TPA and TEN clades consisted only of human isolates. All strains isolated from TNP, including a *Piliocolobus badius* bone derived TPE strain (Tai_105), clustered together in a separate clade within the bigger TPE clade, confirming that syphilis-like lesions and yaws-like lesions observed in this social group of sooty mangabeys were all caused by TPE (Figure 2). Although the MCC tree (Figure 2) depicts a clustering pattern where TNP TPE strains isolated from orofacial lesions appear to have adapted to genital infection and transmission, because they form a well-supported separate internal clade, we argue that this rather supports clustering only by geography, because our additional phylogenetic analyses based on genomes using alternative consensus calling criteria did not support the same clustering pattern (supplementary trees).

**Figure 2:**
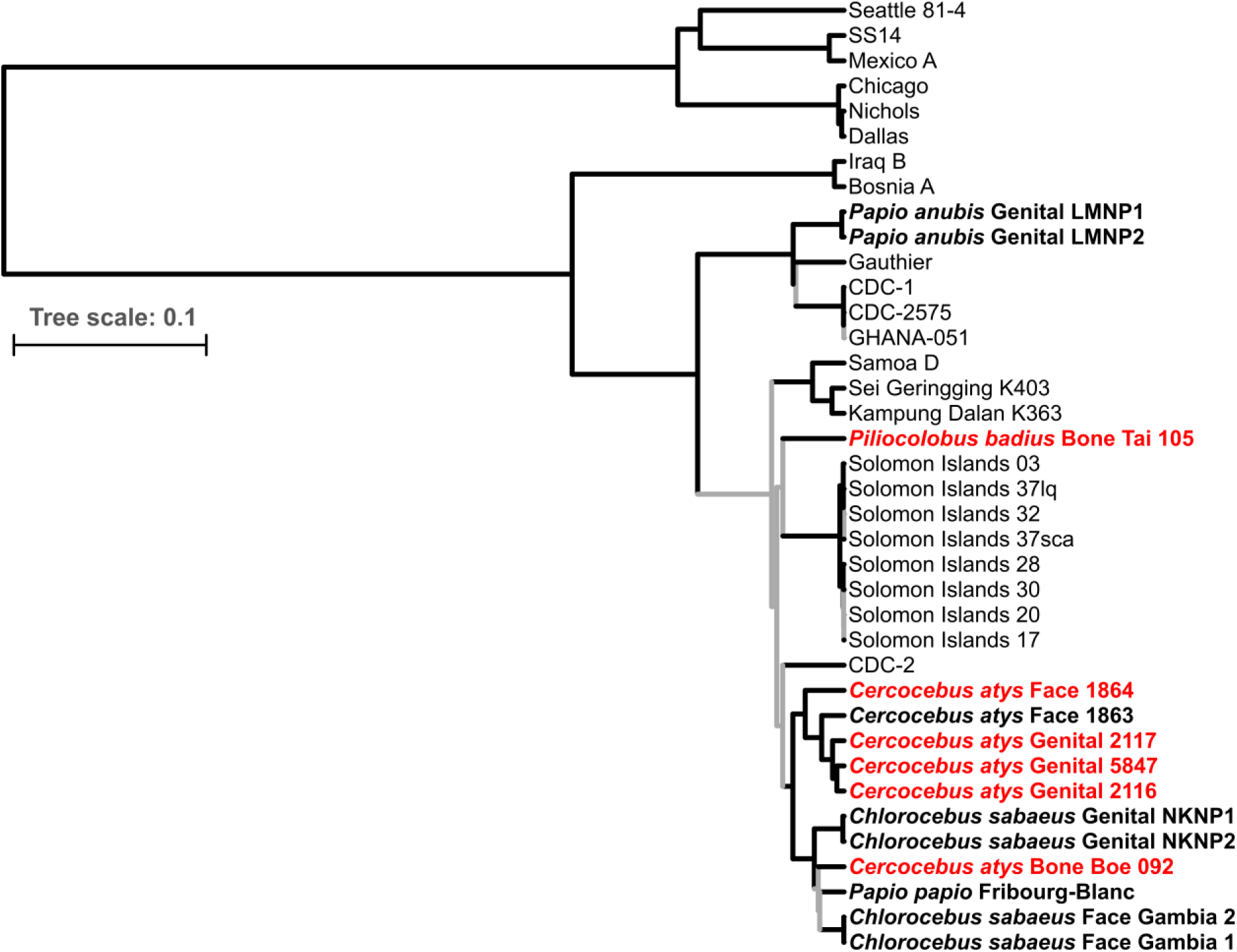
Maximum clade credibility tree of *Treponema pallidum* strains. All simian infecting strains are shown in bold with tip labels showing the species sampled, location of the lesion biopsied or swabbed and sample ID. Genomes generated in this study are shown in red with a minimum coverage of 3X to call a base, and a threshold of 50% identity for a base to be called. The posterior probabilities < 0.95 in the Bayesian Markov chain Monte Carlo tree are indicated in gray. The scale shows nucleotide substitutions per variable site.

In addition, branch lengths (Table S4) showed that strains isolated from TNP exhibited remarkable intra-group evolutionary distance between strains; more than has been observed in TPE infecting any other single population or community of NHP. TNP strains infecting sooty mangabeys had an average reconstructed evolutionary distance of 0.0200% compared to 0.0012% and 0.0000% for the strains infecting green monkeys in Senegal and the Gambia respectively and to 0.0020% for TPE strains infecting olive baboons in Lake Manyara National Park (LMNP, Tanzania). All genomes from TNP were distinct, i.e. we never resampled the same genome despite animals belonging to a single social group. The *Cercocebus atys* bone derived TPE (Boe_092) collected in Guinea Bissau and the Fribourg-Blanc strain isolated from a baboon in Guinea clustered within the clade of TPE strains infecting green monkeys (*Chlorocebus sabaeus*) from the neighbouring Senegal and Gambia. In addition, this green monkey dominated clade formed sister clades with the TNP clade forming a well-supported bigger West African clade. With both phylogenetic inference approaches, simian TPE isolates did not form statistically robust monophyletic groups based on the host species or the types of symptoms animals manifested, but rather clustered according to their geographical origin. Interestingly, simian strains isolated from baboons (LMNP1 and LMNP2) in Tanzania formed a well-supported monophyletic clade together with human infecting TPE strains from Ghana (CDC 1, CDC2, CDC_2575, and Ghana-051).

Bone samples 11786, 22_52, and 15028 had the lowest number of positions covered (0.2% and 62% of genome coverage at 1X). Reads from theses samples were placed onto the final MCC TP phylogeny using the EPA-ng evolutionary placement algorithm [45]. Both chimpanzee (11786 and 15028) and red colobus (*Piliocolobus badius*; 22-52) derived reads from bone samples collected in TNP fell predominantly within the TPE clade of the phylogeny (70% of the reads), with more than 50% of the reads being placed onto the TNP clade. For the red colobus sample 22_52, more than 60% of the reads were placed onto the TPE clade and about 48% were associated with strains from TNP (Figure 3)

**Figure 3:**
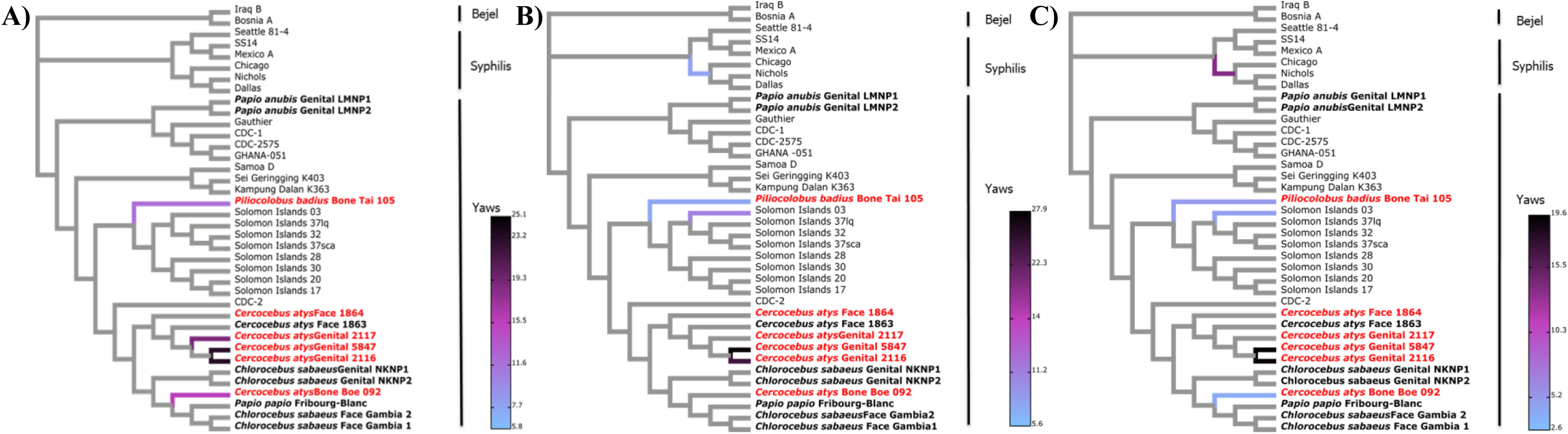
Phylogenetic read placement of bone samples. Heat-trees visualization of phylogenetic placement of TPE mapped reads from bone samples on to the TP MCC (3X overage and 50% threshold) reference tree using the evolutionary placement algorithm (epa-ng). The approximate percentage of reads placed on to a particular branch of the *Treponema pallidum* cladogram is shown as a linearly scaled color density. Genomes generated in this study are shown in red. (A) Sample 11786 (Total number of reads = 517), (B) Sample 15028 (Total number of reads = 3,581), (C) Sample 22_52 (Total number of reads = 19,389)

## Discussion

In the present study, we confirmed the presence of TPE in yaws-like and syphilis-like lesions of sooty mangabeys, joining a growing body of evidence that TPE causes a diversity of symptoms. Yaws-like and syphilis-like lesions caused by TPE infections have been reported in other NHP species [11,28,49–51], however, we present genomic evidence for the occurrence of both symptoms in a single NHP social group inhabiting the same ecosystem. Phylogenetic analyses from reconstructed genomes showed a tree topology where all NHP TP isolates belonged to the TPE clade. However, we found no conclusive evidence to suggest that TPE strains causing the two different pathologies (orofacial and genital lesions) in the sooty mangabey social group formed separate monophyletic groups based on lesions the animals manifested.

Notably, in all TPE strains sampled from this ecosystem, there is no indication that strains were epidemiologically linked, as no identical genomes were sampled. In contrast, in other species in different ecosystems, nearly identical genomes were observed, potentially indicative of an epidemiological link: particularly the two green monkey populations in Bijilo Forest Park (The Gambia) and Niokolo National Park (Senegal) and in baboons from Lake Manyara National Park (Tanzania). The factors driving this high genetic diversity of TPE in TNP are unknown, underscoring the poor understanding of TPE ecology and evolution in wild NHP populations.

In agreement with a recent study of TPE strain diversity among NHPs from Tanzania using multi-locus sequence typing [52], simian derived isolates included in our analyses did not form monophyletic clades based on host species or the type of symptoms caused by an isolate, but rather clustered based on geography. Three geographical clades were prominent; the east African clade consisting of isolates from Tanzania and the two West African sister clades that formed well-supported monophyletic groups. TPE isolates from TNP clustered separately from isolates from NHPs in far West Africa, particularly Guinea, Guinea Bissau, Senegal and The Gambia, which are all close neighbouring countries (Figure 2). The far West Africa clade consisted of TPE strains isolated from green monkeys, guinea baboon, and a bone derived sequence from a sooty mangabey, all from ecosystems in close proximity to one another. The formation of these sister clades irrespective of the symptoms observed or the host species from which the strains were derived suggests that geography and the ecosystems NHPs inhabit may be important factors influencing the diversification of NHP TPE strains [53]. The phylogenetic read placement of TPE reads derived from chimpanzee (11786 and 15028) and red colobus (22-52) bone samples from TNP (Figure 3) onto the TNP sooty mangabey clade provides further support to the suggestion that geography shapes the phylogenetic relationships among simian strains. Such a clustering pattern might indicate cross-species transmission of yaws between species sharing the same habitat or infection of NHP sharing a habitat from some shared unknown source [17, 54].

One transmission mode that has been suggested is vectorial transmission. Under experimental conditions, viable TP spirochetes were transmitted by flies between different host species causing clinical disease [55, 56]. Recently, Gogarten *et al*. (2019) showed that flies carrying the yaws pathogen formed high-density persistent associations with NHP social groups in the TNP. These stable high-density fly-NHP associations might provide opportunities for TPE transmission by flies [57]. Knauf *et al*. (2016) also isolated TP DNA from necrophagous flies in ecosystems where TP infections in NHPs are common [58]. Further work is needed to confirm whether flies can actually transmit TPE in the wild [59].

Interspecies interactions that could facilitate transmission via direct contact between NHP species inhabiting TNP are well-documented; these include a strong predator-prey relationship between chimpanzees and red colobus monkeys, while the overlapping home ranges and large amounts of time spent in mixed-species associations provides opportunities for between-species transmission through various kinds of direct contact (e.g., grooming, fighting, play, mating) [60, 61]. A sexual transmission mode has been suggested due to the predilection of genital lesions in individuals of reproductive age [62], though younger animals are also infected in the TNP mangabey group. Further research is also needed to understand the transmission of TPE between NHP.

We detected support for a clade comprising human infecting TPE strains from Ghana and NHP infecting TPE strains from Tanzania. While intriguing, considering the wide spatio-temporal relationships of these strains, we do not consider this to be definitive evidence for cross-species transmission. Indeed, genomic data from human infections is largely lacking from these countries. For example, to date no human strains from Côte d’Ivoire have been sequenced, despite ongoing human infections [12]. Given the spatial signal among simian isolates detected here, future studies should investigate human yaws infections in these regions and determine whether such a spatial signal also exists for human-infecting yaws-causing bacteria. Such data will help to determine whether inter-species transmission of TPE between humans and NHPs occurs, particularly as the sampling of NHP TPE genetic diversity has greatly improved over the last several years. In fact, of the thirty TPE genomes included in the current phylogenomic analyses, fourteen originate from NHP.

This study joins a growing body of evidence that human and wildlife bones may contain a treasure trove of data on treponemes [21, 63]. We detected TPE DNA in NHP bones from TNP and four additional sites across sub-Saharan Africa (Table S2) and reconstructed partial genomes from several TPE positive bones. The finding of TPE in bones confirms that NHP infections existed in the park for at least three decades (Table S2), complementing clinical evidence which only started accumulating from 2014 on [12]. In addition, despite the lack of clinical evidence, the successful phylogenetic placement of bone derived TPE reads derived from chimpanzee and red colobus monkey bones onto the TNP clade, suggests that sooty mangabeys are not the only NHP species affected in the park and lends support to local transmission of TPE within ecosystems. Finally, the detection of TPE DNA in bones from sub-Saharan Africa sites where no clinical cases have been reported in NHPs, suggests that TPE circulation in NHPs is underreported. In this regard, increased surveillance of TPE in wildlife is needed and will expand our knowledge of the ecological niche of this pathogen and could be useful for informing eradication efforts. Despite the knowledge that TPE has been infecting NHPs since at least the late 1960s [50, 51], only a single isolate dating back to this period is available [13]. Future studies extracting genomic data from old NHP bones or human medical archival materials from yaws endemic regions may provide important insights into TPE ecology and evolution.

## Conclusions

There is a large diversity of TPE strains infecting NHPs in TNP, even within a single social group. Phylogenetic patterns observed in this study are compatible with cross-species transmission of TPE within ecosystems, though how often and by what means this transmission occurs remains an important area of future research.

## Data availability

We archived all raw sequence read files in NCBI under BioProject PRJNA588802.

## Acknowledgements

This article represents a chapter in the doctoral dissertation of B.M. whose research activities were financed by the Robert Koch Institute, Berlin, Germany. We thank the Ivorian directorship of the TaïNP, the Office Ivoirien des Parcs et Réserves, the Centre Suisse de Recherche Scientifique, the Taï Chimpanzee Project and the Taï Monkey project and their teams of field assistants for their support. We thank the following people for their support with field and organizational activities; Christophe Boesch, Katherine Corogenes, Karsten Dierks, Dervla Dowd, Henk Eshuis, Annemarie Goedmakers, John Hart, Thurston Cleveland Hicks, Theo Freeman, Sorrel Jones, Vincent Lapeyre, Juan Lapuente, Vera Leinert, Sergio Marrocoli, Amelia Meier, Yasmin Moebius, Geoffrey Muhanguzi, Mizuki Murai, Emmanuelle Normand, Martha M. Robbins, Joost van Schijndel, Volker Sommer, Virginie Vergnes, and Klaus Zuberbuehler. For their funding support, we thank the Max Planck Society, Max Planck Society Innovation Fund, and Heinz L. Krekeler Foundation. We are also grateful to the following institutions and Ministries in the following countries for facilitating activities pertaining to this study: Ministere des Eaux et Forets, Côte d’Ivoire; Institut Congolais pour la Conservation de la Nature, DR-Congo; Ministere de la Recherche Scientifique, DR-Congo; Agence Nationale des Parcs Nationaux, Gabon; Centre National de la Recherche Scientifique (CENAREST), Gabon: Société Equatoriale d’Exploitation Forestière (SEEF), Gabon; Ministere de l’Agriculture de l’Elevage et des Eaux et Forets, Guinea; Instituto da Biodiversidade e das Áreas Protegidas (IBAP); Ministro da Agricultura e Desenvolvimento Rural, Guinea-Bissau; Forestry Development Authority, Liberia; Conservation Society of Mbe Mountains (CAMM), Nigeria; National Park Service, Nigeria; Direction des Eaux, Forêts et Chasses, Senegal; Makerere University Biological Field Station (MUBFS), Uganda; Uganda National Council for Science and Technology (UNCST), Uganda. We also thank the following non-governmental organisations for the their assistance and support of field activities: Wild Chimpanzee Foundation, Côte d’Ivoire; Lukuru Wildlife Research Foundation, DRC; WCS Albertine Rift Programme, DRC; WWF Congo Basin, DRC; Loango Ape Project, Gabon; The Aspinall Foundation, Gabon; Station d’Etudes des Gorilles et Chimpanzees, Gabon: Kwame Nkrumah University of Science and Technology (KNUST), Ghana; Wild Chimpanzee Foundation, Guinea; Foundation Chimbo (Boe); Wild Chimpanzee Foundation, Liberia; Gashaka Primate Project, Nigeria; Wildlife Conservation Society (WCS) Nigeria, Nigeria; Fongoli Savanna Chimpanzee Project, Senegal; Jane Goodall Insitute Spain (Dindefelo), Senegal; Budongo Conservation Field Station, Uganda; Ngogo Chimpanzee Project, Uganda.

---

**Table S1:**
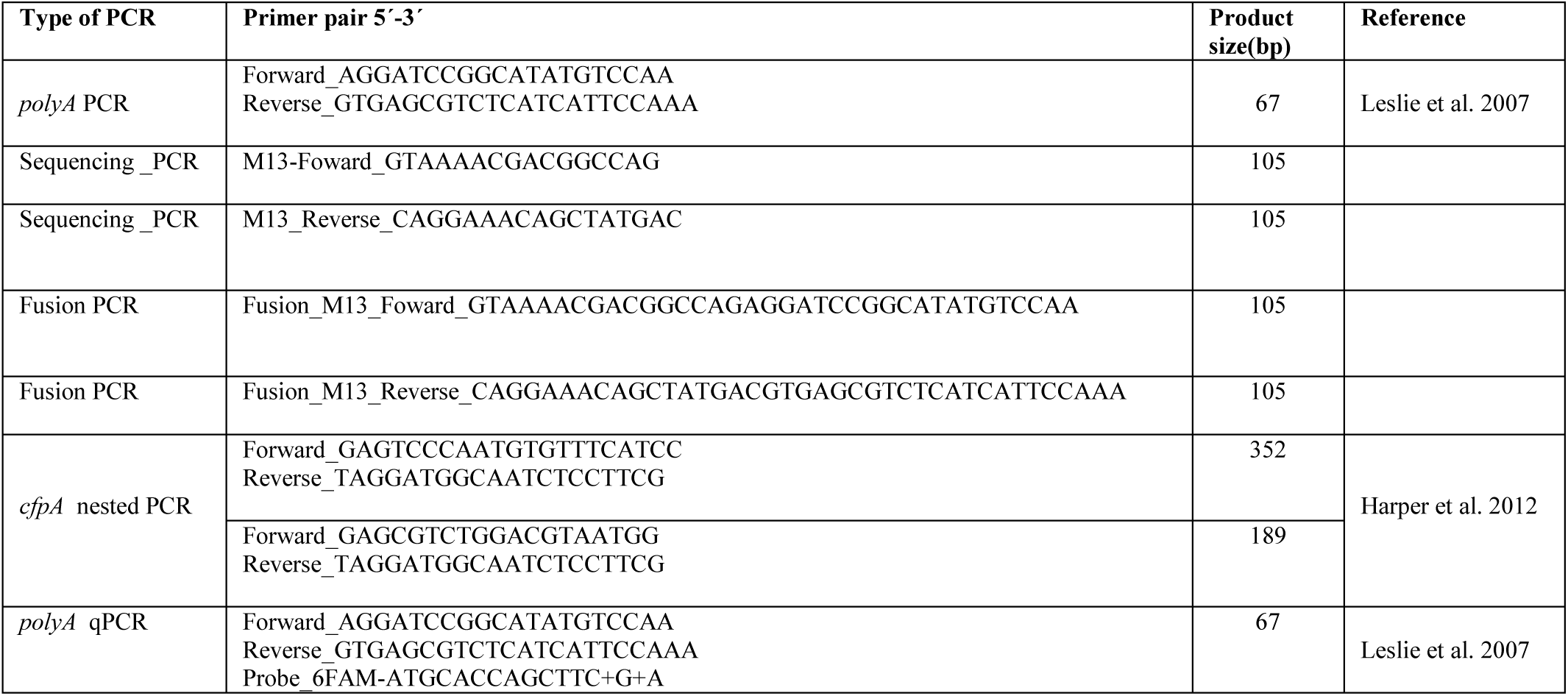
Primers used for the Treponema pallidum screening.

**Table S2: List of bone sample used in this study (see supplementary material 1)**

**Table S3:** Published genomes used in this study.

| Isolate ID | RefSeq Accession ID | Host | TP spectrum |
| --- | --- | --- | --- |
| Bosnia A | CP007548.1 | *Homo Sapiens* | Bejel |
| Iraq_B | CP032303.1 | *Homo Sapiens* | Bejel |
| Nichols | NC_021490.2 | *Homo Sapiens* | Syphilis |
| SS14 | NC_021508.1 | *Homo Sapiens* | Syphilis |
| Chicago | NC_017268.1 | *Homo Sapiens* | Syphilis |
| Mexico A | NC_018722.1 | *Homo Sapiens* | Syphilis |
| Dallas | NC_016844.1 | *Homo Sapiens* | Syphilis |
| Seattle 81-4 | CP003679.1 | *Homo Sapiens* | Syphilis |
| Fribourg-Blanc | NC_021179.1 | *Papio cynocephalus* | Yaws |
| Samoa D | NC_016842.1 | *Homo Sapiens* | Yaws |
| Gauthier | NC_016843.1 | *Homo Sapiens* | Yaws |
| CDC-1 | CP024750.1 | *Homo Sapiens* | Yaws |
| CDC-2 | NC_016848.1 | *Homo Sapiens* | Yaws |
| CDC_2575 | CP020366 | *Homo Sapiens* | Yaws |
| Ghana-051 | CP020365 | *Homo Sapiens* | Yaws |
| Kampung_Dalan_K363 | CP024088.1 | *Homo Sapiens* | Yaws |
| Sei_Geringging_K403 | CP024089.1 | *Homo Sapiens* | Yaws |
| Solomon Islands 03 | ERR1470343 | *Homo Sapiens* | Yaws |
| Solomon Islands 17 | ERR1470344 | *Homo Sapiens* | Yaws |
| Solomon Islands 20 | ERR1470335 | *Homo Sapiens* | Yaws |
| Solomon Islands 28 | ERR1470338 | *Homo Sapiens* | Yaws |
| Solomon Islands 30 | ERR1470334 | *Homo Sapiens* | Yaws |
| Solomon Islands 32 | ERR1470342 | *Homo Sapiens* | Yaws |
| Solomon Islands 37 liq | ERR1470330 | *Homo Sapiens* | Yaws |
| Solomon Islands 37 sca | ERR1470331 | *Homo Sapiens* | Yaws |
| Gambia-1 | SRR4308597 | *Chlorocebus sabeus* | Yaws |
| Gambia-2 | SRR4308605 | *Chlorocebus sabaeus* | Yaws |
| Senegal NKNP-1 | SRR4308606 | *Chlorocebus sabaeus* | Yaws |
| Senegal NKNP-2 | SRR4308607 | *Chlorocebus sabaeus* | Yaws |
| LMNP-1 | CP021113.1 | *Papio anubis* | Yaws |
| LMNP-2_BS5 | SRR4308598 | *Papio anubis* | Yaws |
| LMNP-2_BS6 | SRR4308599 | *Papio anubis* | Yaws |
| LMNP-2_BS7 | SRR4308601 | *Papio anubis* | Yaws |
| LMNP-2_BS8 | SRR4308602 | *Papio anubis* | Yaws |
| 1863-Hato | SRR4308604 | *Cercocebus atys* | Yaws |

**Table S4: Pairwise patristic distance of ML tree (TPE and TEN strains only) –See supplementary material 2**

## Supplementary trees

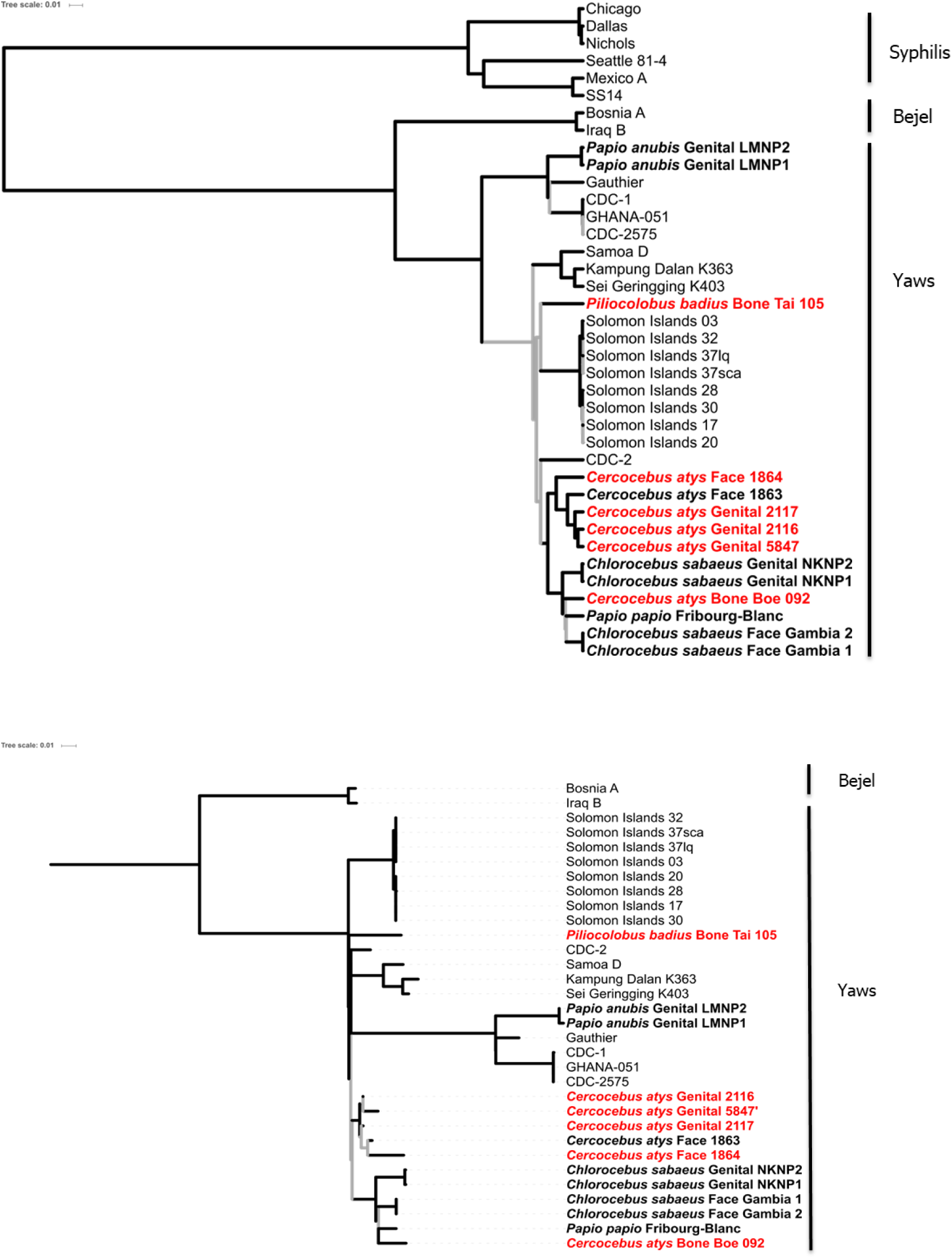

**Maximum clade credibility and maximum likelihood trees at 3X coverage and 95% threshold**; Maximum likelihood tree shown is for TPE and TEN intra-group phylogenomic anylsis only. Branch support was assessed using Shimodaira-Hasegawa-like approximate likelihood ratio tests (SH-like aLRT), with branches supported by SH-like aLRT values < 0.90 and/or posterior probabilities < 0.95 in the Bayesian Markov chain Monte Carlo tree indicated in gray. The scale shows nucleotide substitutions per variable site.

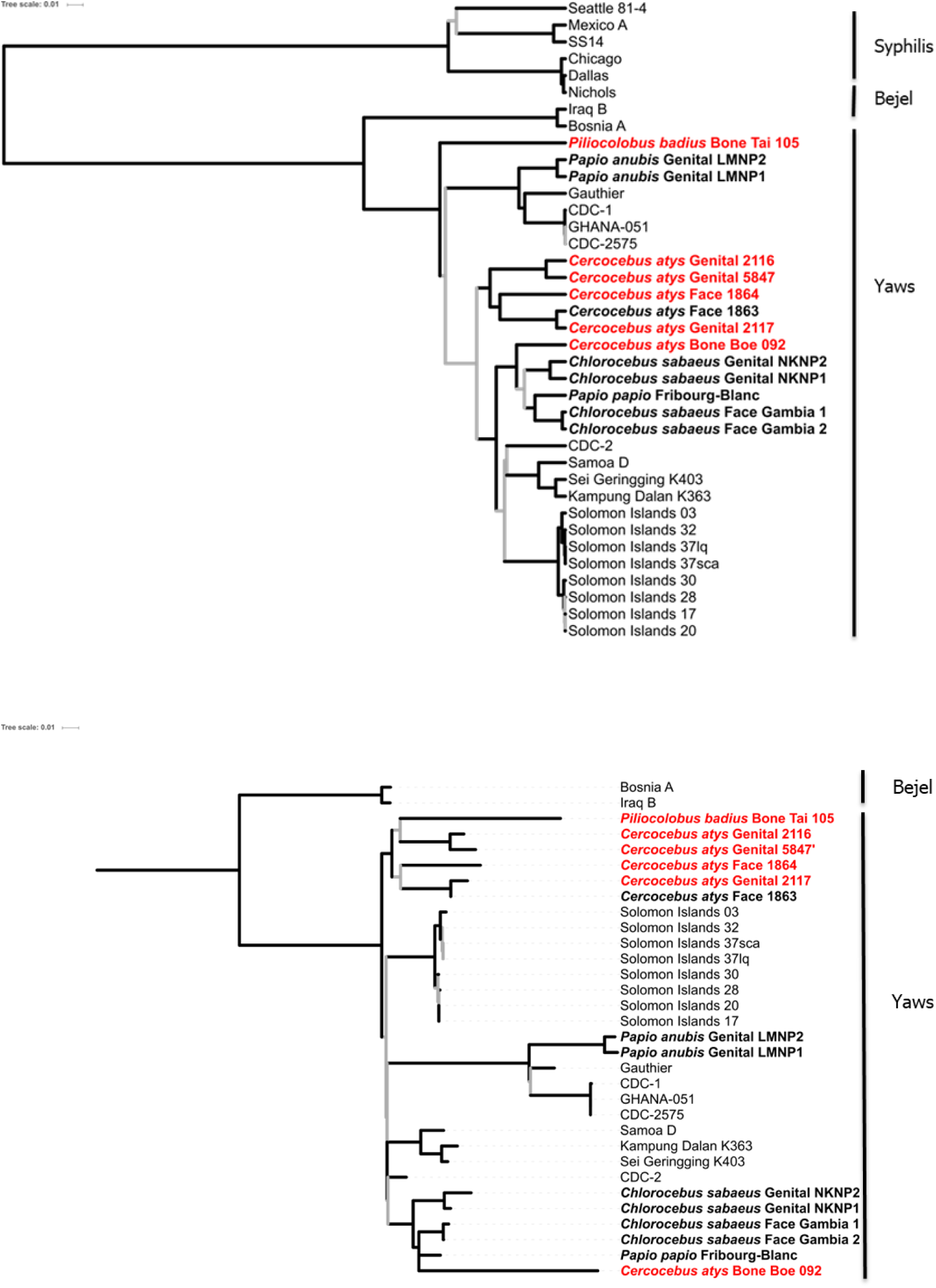

**Maximum clade credibility and maximum likelihood trees at 5X coverage and 50% threshold**; Maximum likelihood tree shown is for TPE and TEN intra-group phylogenomic anylsis only. Branch support was assessed using Shimodaira-Hasegawa-like approximate likelihood ratio tests (SH-like aLRT), with branches supported by SH-like aLRT values < 0.90 and/or posterior probabilities < 0.95 in the Bayesian Markov chain Monte Carlo tree indicated in gray. The scale shows nucleotide substitutions per variable site.

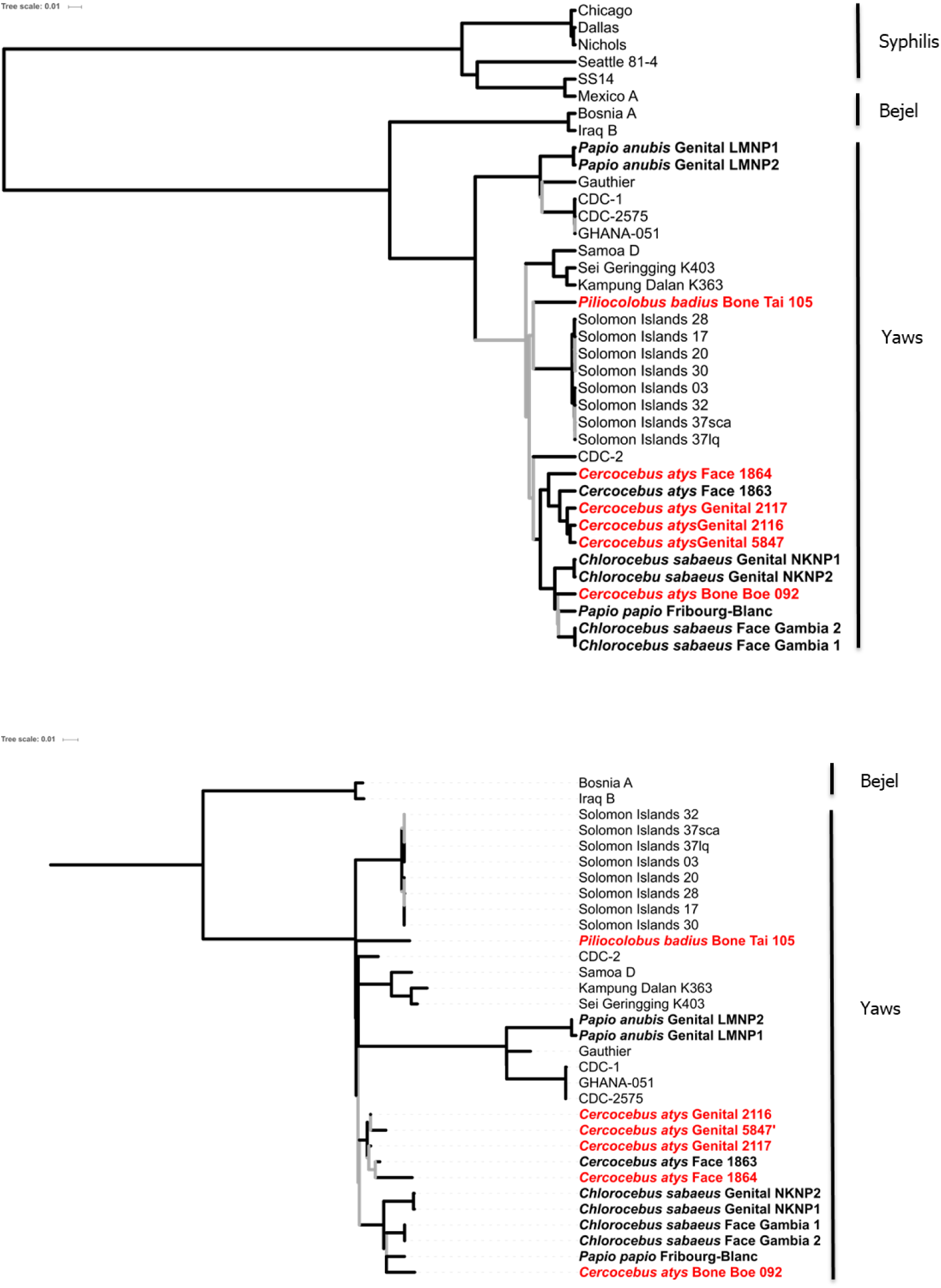

**Maximum clade credibility and maximum likelihood trees at 5X coverage and 95% threshold**; Maximum likelihood tree shown is for TPE and TEN intra-group phylogenomic anylsis only. Branch support was assessed using Shimodaira-Hasegawa-like approximate likelihood ratio tests (SH-like aLRT), with branches supported by SH-like aLRT values < 0.90 and/or posterior probabilities < 0.95 in the Bayesian Markov chain Monte Carlo tree indicated in gray. The scale shows nucleotide substitutions per variable site.

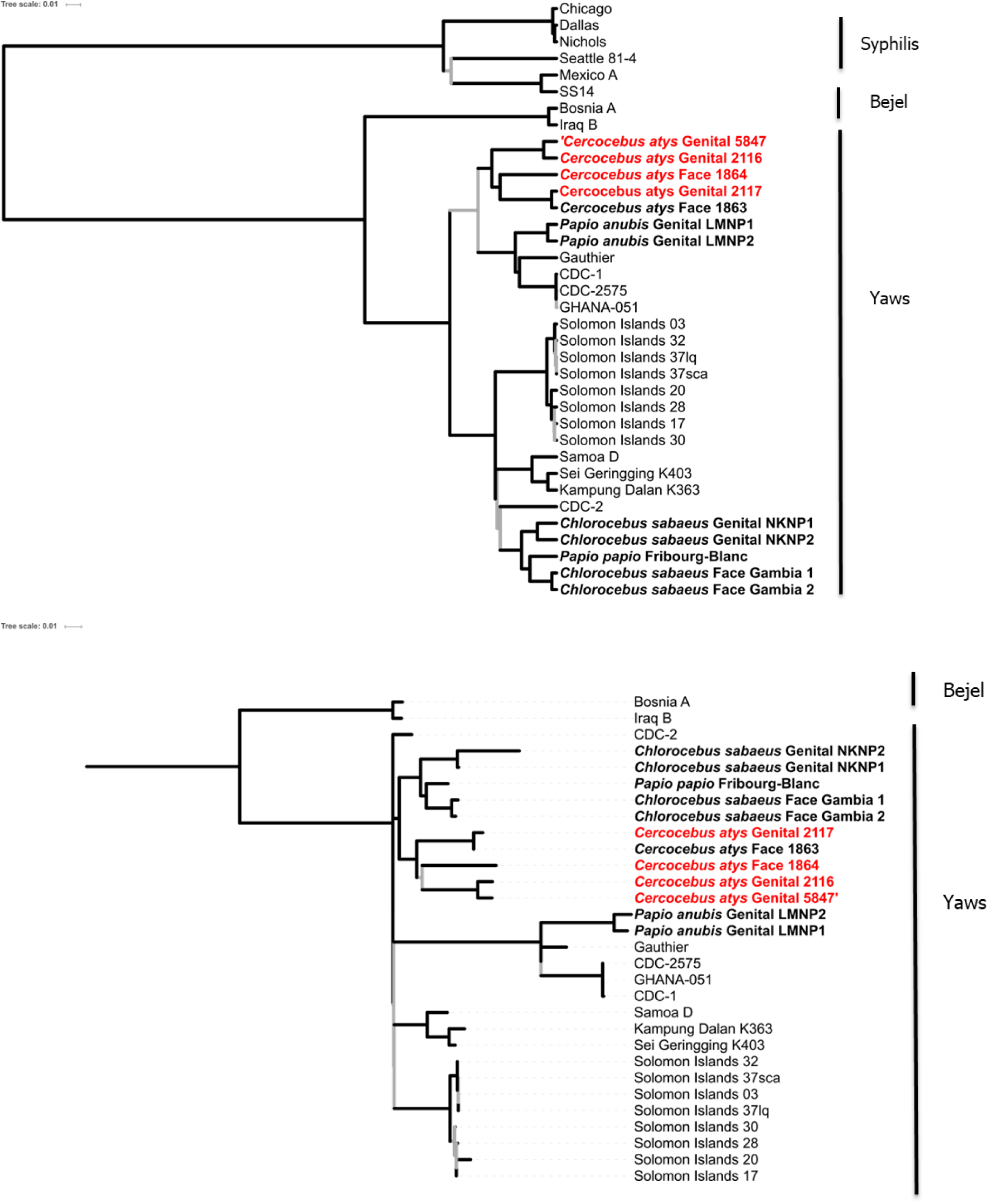

**Maximum clade credibility and maximum likelihood trees at 10X coverage and 50% threshold**; Maximum likelihood tree shown is for TPE and TEN intra-group phylogenomic anylsis only. Branch support was assessed using Shimodaira-Hasegawa-like approximate likelihood ratio tests (SH-like aLRT), with branches supported by SH-like aLRT values < 0.90 and/or posterior probabilities < 0.95 in the Bayesian Markov chain Monte Carlo tree indicated in gray. The scale shows nucleotide substitutions per variable site.

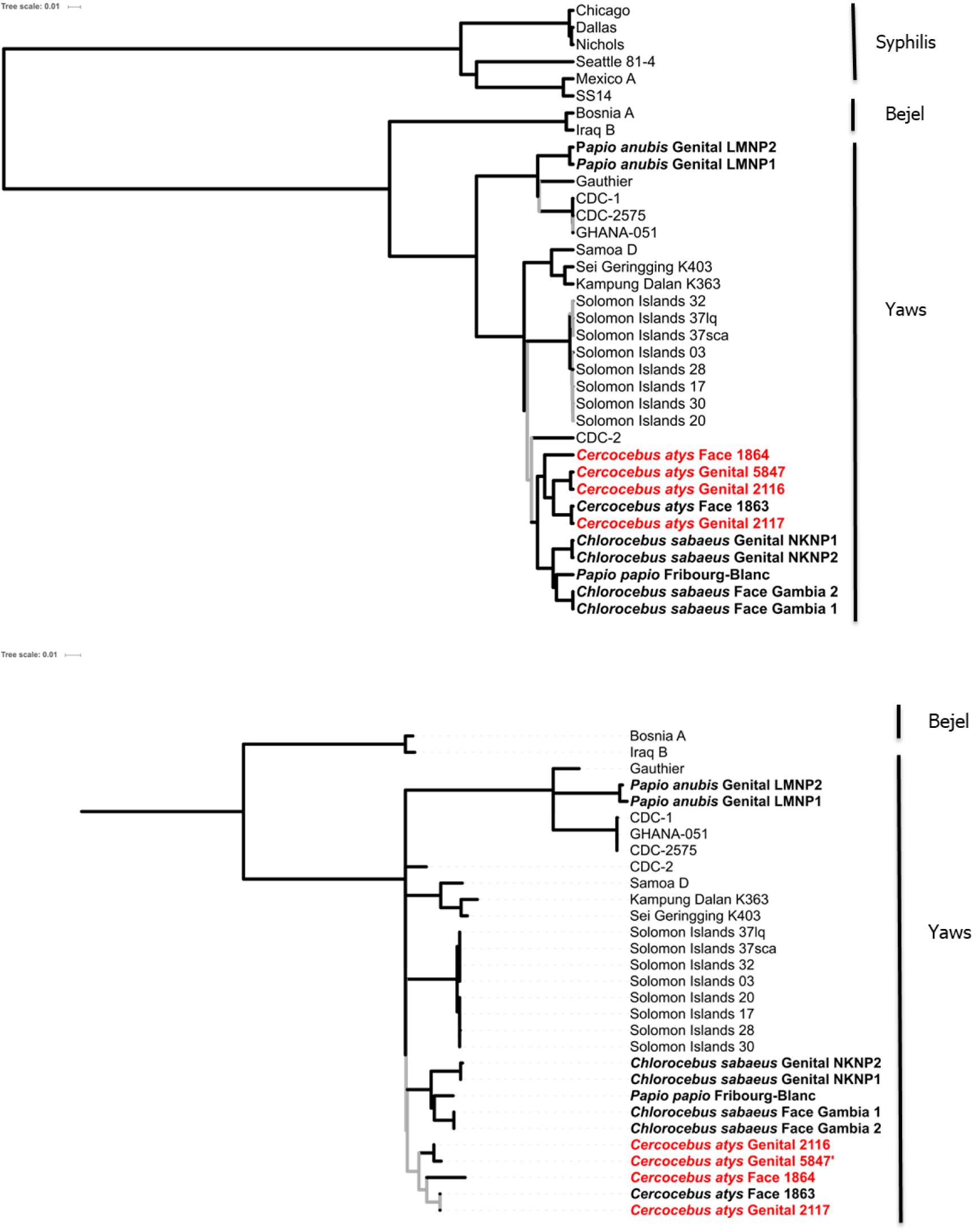

**Maximum clade credibility and maximum likelihood trees at 10X coverage and 95% threshold**; Maximum likelihood tree shown is for TPE and TEN intra-group phylogenomic anylsis only. Branch support was assessed using Shimodaira-Hasegawa-like approximate likelihood ratio tests (SH-like aLRT), with branches supported by SH-like aLRT values < 0.90 and/or posterior probabilities < 0.95 in the Bayesian Markov chain Monte Carlo tree indicated in gray. The scale shows nucleotide substitutions per variable site.

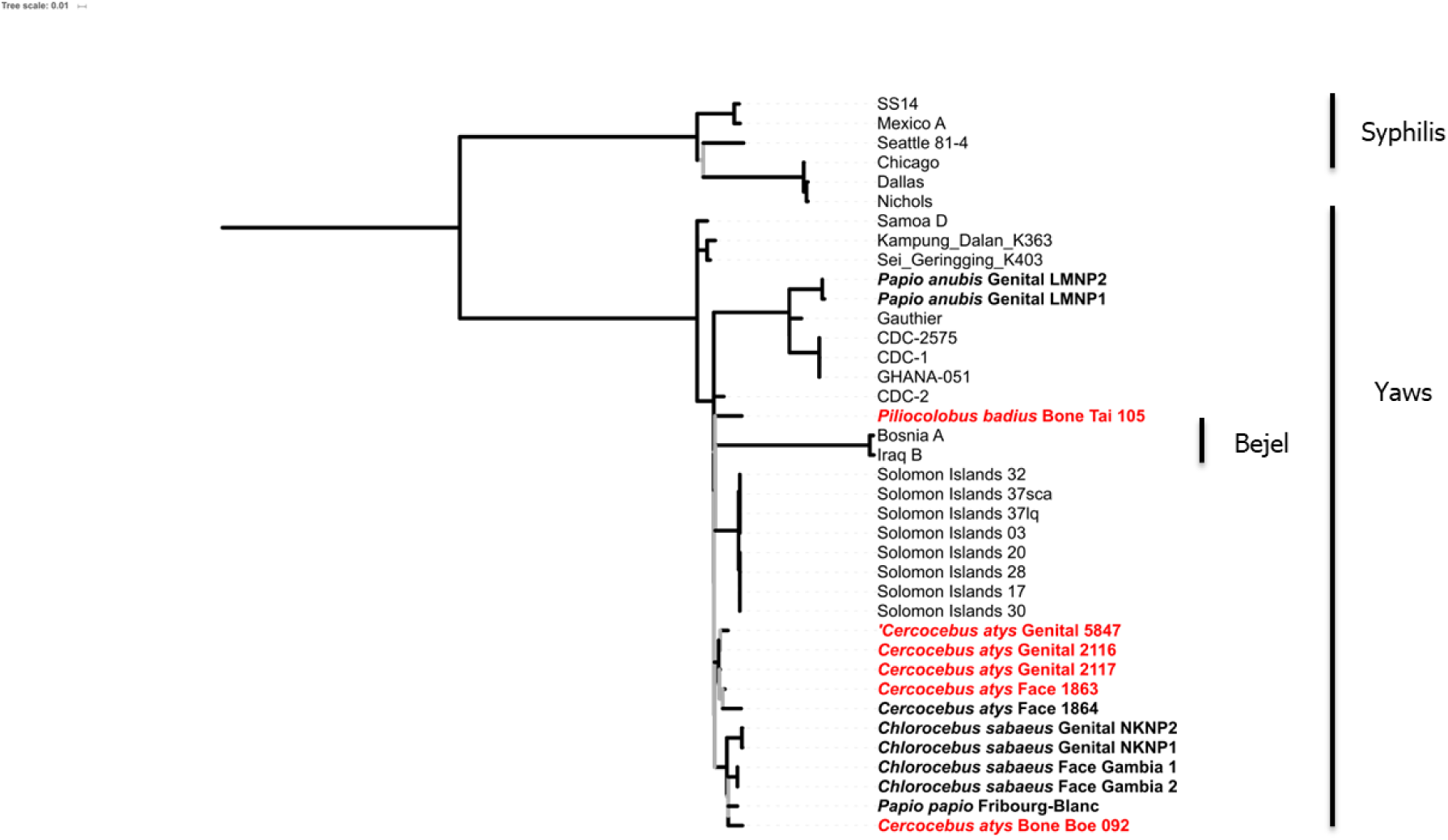

**Maximum likelihood tree at 3X coverage and 50% threshold**; shown here is the TEN clade clustering within the TPE clade. This clustering pattern was the same for all ML trees and hence the control intra-group analysis of TPE and TEN only. Branch support was assessed using Shimodaira-Hasegawa-like approximate likelihood ratio tests (SH-like aLRT), with branches supported by SH-like aLRT values < 0.90 indicated in gray. The scale shows nucleotide substitutions per variable site.

## References

1. Armelagos GJ, Zuckerman MK, Harper KN. The science behind pre-columbian evidence of syphilis in europe: research by documentary. Evol Anthropol Issues, News, Rev. 2012;21: 50–57. doi:10.1002/evan.20340

2. Lithgow K V, Hof R, Wetherell C, Phillips D, Houston S, Cameron CE. A defined syphilis vaccine candidate inhibits pallidum. Nat Commun. 2017;8: 1–10. doi:10.1038/ncomms14273

3. Giacani L, Lukehart SA. The endemic treponematoses. Clin Microbiol Rev. 2014;27: 89–115. doi:10.1128/CMR.00070-13

4. Marks M, Solomon AW, Mabey DC. Endemic treponemal diseases. Trans R Soc Trop Med Hyg. 2014;108: 601–7. doi:10.1093/trstmh/tru128

5. Čejková D, Zobaníková M, Chen L, Pospíšilová P, Strouhal M, Qin X, et al. Whole genome sequences of three *Treponema pallidum* ssp. *pertenue* strains: Yaws and syphilis treponemes differ in less than 0.2% of the genome sequence. PLoS Negl Trop Dis. 2012;6. doi:10.1371/journal.pntd.0001471

6. Centurion-Lara A, Molini BJ, Godornes C, Sun E, Hevner K, Van Voorhis WC, et al. Molecular differentiation of *Treponema pallidum* subspecies. J Clin Microbiol. 2006;44: 3377–80. doi:10.1128/JCM.00784-06

7. Newman L, Rowley J, Vander Hoorn S, Wijesooriya NS, Unemo M, Low N, et al. Global estimates of the prevalence and incidence of four curable sexually transmitted infections in 2012 based on systematic review and global reporting. Meng Z, editor. PLoS One. 2015;10: e0143304. doi:10.1371/journal.pone.0143304

8. World Health Organisation. First WHO report on neglected tropical diseases: working to overcome the global impact of neglected tropical diseases. [Internet]. 2010. doi:WHO/HTM/NTD/2010.1

9. World Health Organisation. Global health sector strategy on sexually transmitted infections, 2016-2021 [Internet]. 2016. Report No.: WHO/RHR/16.09. Available: https://web.archive.org/web/20181019130311/http://www.who.int/reproductivehealth/publications/rtis/ghss-stis/en/

10. Dyson L, Mooring EQ, Holmes A, Tildesley MJ, Marks M. Insights from quantitative and mathematical modelling on the proposed 2030 goals for yaws. Gates Open Res. 2019;3: 1576. doi:10.12688/gatesopenres.13078.1

11. Chuma IS, Batamuzi EK, Collins DA, Fyumagwa RD, Hallmaier-Wacker LK, Kazwala RR, et al. Widespread *Treponema pallidum* infection in nonhuman primates, Tanzania. Emerg Infect Dis. 2018;24: 1002–1009. doi:10.3201/eid2406.180037

12. Knauf S, Gogarten JF, Schuenemann VJ, Nys HM De, Düx A, Strouhal M, et al. Nonhuman primates across sub-Saharan Africa are infected with the yaws bacterium *Treponema pallidum* subsp. *pertenue*. Emerg Microbes Infect. 2018; doi:10.1038/s41426-018-0156-4

13. Zobaníková M, Strouhal M, Mikalová L, Čejková D, Ambrožová L, Pospíšilová P, et al. Whole genome sequence of the treponema Fribourg-blanc: unspecified simian isolate is highly similar to the yaws subspecies. PLoS Negl Trop Dis. 2013;7: e2172. doi:10.1371/journal.pntd.0002172

14. Smith JL, David NJ, Indgin S, Israel CW, Levine BM, Justice J, et al. Neuro-ophthalmological study of late yaws and pinta. II. The Caracas project. Sex Transm Infect. 1971;47: 226–251. doi:10.1136/sti.47.4.226

15. Lawton Smith J, Israel CW. Recovery of spirochagetes in the monkey by passive transfer from human late sero-negative syphilis. Br J Vener Dis. 1968;44: 109–15. doi:10.1136/sti.44.2.109

16. Plowright RK, Parrish CR, McCallum H, Hudson PJ, Ko AI, Graham AL, et al. Pathways to zoonotic spillover. Nat Rev Microbiol. 2017;15: 502.

17. Pedersen AB, Davies TJ. Cross-species pathogen transmission and disease emergence in primates. Ecohealth. 2009;6: 496–508. doi:10.1007/s10393-010-0284-3

18. Han BA, Kramer AM, Drake JM. Global patterns of zoonotic disease in mammals. Trends Parasitol. 2016;32: 565–577.

19. Geoghegan JL, Duchêne S, Holmes EC. Comparative analysis estimates the relative frequencies of co-divergence and cross-species transmission within viral families. Drosten C, editor. PLOS Pathog. 2017;13: e1006215. doi:10.1371/journal.ppat.1006215

20. Gao F, Bailes E, Robertson DL, Chen Y, Rodenburg CM, Michael SF, et al. Origin of HIV-1 in the chimpanzee *Pan troglodytes troglodytes*. Nature. 1999;397: 436–441. doi:10.1038/17130

21. Gogarten JF, Düx A, Schuenemann VJ, Nowak K, Boesch C, Wittig RM, et al. Tools for opening new chapters in the book of *Treponema pallidum* evolutionary history. Clin Microbiol Infect. 2016;22: 916–921. doi:10.1016/j.cmi.2016.07.027

22. Calvignac-Spencer S, Merkel K, Kutzner N, Kühl H, Boesch C, Kappeler PM, et al. Carrion fly-derived DNA as a tool for comprehensive and cost-effective assessment of mammalian biodiversity. Mol Ecol. 2013;22: 915–924. doi:10.1111/mec.12183

23. Hoffmann C, Zimmermann F, Biek R, Kuehl H, Nowak K, Mundry R, et al. Persistent anthrax as a major driver of wildlife mortality in a tropical rainforest. Nature. 2017;548: 82–86. doi:10.1038/nature23309

24. Orlando L, Ginolhac A, Raghavan M, Vilstrup J, Rasmussen M, Magnussen K, et al. True single-molecule DNA sequencing of a pleistocene horse bone. Genome Res. 2011;21: 1705–1719. doi:10.1101/gr.122747.111

25. Rohland N, Hofreiter M. Ancient DNA extraction from bones and teeth. Nat Protoc. 2007;2: 1756–1762.

26. Gamba C, Hanghøj K, Gaunitz C, Alfarhan AH, Alquraishi SA, Al-Rasheid KAS, et al. Comparing the performance of three ancient DNA extraction methods for high-throughput sequencing. Mol Ecol Resour. 2015;

27. Leslie DE, Azzato F, Karapanagiotidis T, Leydon J, Fyfe J. Development of a real-time PCR assay to detect *Treponema pallidum* in clinical specimens and assessment of the assay’s performance by comparison with serological testing. J Clin Microbiol. 2007;45: 93–96. doi:10.1128/JCM.01578-06

28. Harper KN, Fyumagwa RD, Hoare R, Wambura PN, Coppenhaver DH, Sapolsky RM, et al. *Treponema pallidum* infection in the wild baboons of east africa: distribution and genetic characterization of the strains responsible. PLoS One. 2012;7. doi:10.1371/journal.pone.0050882

29. Altschul SF, Gish W, Miller W, Myers EW, Lipman DJ. Basic local alignment search tool. J Mol Biol. 1990;215: 403–410. doi:10.1016/S0022-2836(05)80360-2

30. Bolger AM, Lohse M, Usadel B. Trimmomatic: a flexible trimmer for Illumina sequence data. Bioinformatics. 2014;30: 2114–2120. Available: https://www.ncbi.nlm.nih.gov/pmc/articles/PMC4103590/pdf/btu170.pdf

31. Peltzer A, Jäger G, Herbig A, Seitz A, Kniep C, Krause J, et al. EAGER: efficient ancient genome reconstruction. Genome Biol. 2016;17: 60.

32. Li H, Handsaker B, Wysoker A, Fennell T, Ruan J, Homer N, et al. The sequence alignment/map format and SAMtools. Bioinformatics. 2009;25: 2078–2079.

33. Kearse M, Moir R, Wilson A, Stones-Havas S, Cheung M, Sturrock S, et al. Geneious Basic: An integrated and extendable desktop software platform for the organization and analysis of sequence data. Bioinformatics. 2012;28: 1647–1649. doi:10.1093/bioinformatics/bts199

34. Katoh K, Standley DM. MAFFT multiple sequence alignment software version 7: improvements in performance and usability. Mol Biol Evol. 2013;30: 772–80. doi:10.1093/molbev/mst010

35. Arora N, Schuenemann VJ, Jäger G, Peltzer A, Seitz A, Herbig A, et al. Origin of modern syphilis and emergence of a pandemic *Treponema pallidum* cluster. Nat Microbiol. 2017;2: 16245. doi:10.1038/nmicrobiol.2016.245

36. Talavera G, Castresana J. Improvement of phylogenies after removing divergent and ambiguously aligned blocks from protein sequence alignments. Kjer K, Page R, Sullivan J, editors. Syst Biol. 2007;56: 564–577. doi:10.1080/10635150701472164

37. Gouy M, Guindon S, Gascuel O. SeaView Version 4: A Multiplatform graphical user interface for sequence alignment and phylogenetic tree building. Mol Biol Evol. 2010;27: 221–224. doi:10.1093/molbev/msp259

38. Lefort V, Longueville J-E, Gascuel O. SMS: Smart model selection in PhyML. Mol Biol Evol. 2017;34: 2422–2424. doi:10.1093/molbev/msx149

39. Guindon S, Dufayard J-F, Lefort V, Anisimova M, Hordijk W, Gascuel O. New algorithms and methods to estimate maximum-likelihood phylogenies: assessing the performance of PhyML 3.0. Syst Biol. 2010;59: 307–321. doi:10.1093/sysbio/syq010

40. Rambaut A, Lam TT, Carvalho LM, Pybus OG. Exploring the temporal structure of heterochronous sequences using TempEst (formerly Path-O-Gen). Virus Evol. 2016;2: vew007. doi:10.1093/VE/VEW007

41. Fourment M, Gibbs M. PATRISTIC: a program for calculating patristic distances and graphically comparing the components of genetic change. BMC Evol Biol. 2006;6: 1. doi:10.1186/1471-2148-6-1

42. Rambaut A, Drummond AJ, Xie D, Baele G, Suchard MA. Posterior summarization in Bayesian phylogenetics using Tracer 1.7. Syst Biol. 2018;67: 901–904.

43. Drummond AJ, Rambaut A. BEAST: Bayesian evolutionary analysis by sampling trees. BMC Evol Biol. 2007;7: 214.

44. Letunic I, Bork P. Interactive tree of life (iTOL) v4: recent updates and new developments. Nucleic Acids Res. 2019; doi:10.1093/nar/gkz239

45. Barbera P, Kozlov AM, Czech L, Morel B, Darriba D, Flouri T, et al. EPA-ng: Massively parallel evolutionary placement of genetic sequences. Posada D, editor. Syst Biol. 2019;68: 365–369. doi:10.1093/sysbio/syy054

46. Berger SA, Stamatakis A. Aligning short reads to reference alignments and trees. Bioinformatics. 2011;27: 2068–2075. doi:10.1093/bioinformatics/btr320

47. Kozlov AM, Darriba D, Flouri T, Morel B, Stamatakis A. RAxML-NG: a fast, scalable and user-friendly tool for maximum likelihood phylogenetic inference. Wren J, editor. Bioinformatics. 2019; doi:10.1093/bioinformatics/btz305

48. Czech L, Barbera P, Stamatakis A. Genesis and Gappa: library and toolkit for working with phylogenetic (placement) data. bioRxiv. 2019; 647958. doi:10.1101/647958

49. Knauf S, Batamuzi EK, Mlengeya T, Kilewo M, Lejora IAV, Nordhoff M, et al. Treponema infection associated with genital ulceration in wild baboons. Vet Pathol. 2012;49: 292–303. doi:10.1177/0300985811402839

50. Baylet R, Thivolet J, Sepetjian M, Nouhouay Y, Baylet M. [Natural open treponematosis in the *Papio papio* baboon in Casamance]. Bull Soc Pathol Exot Filiales. 1971;64: 842–846.

51. Fribourg-Blanc A, Mollaret HH. Natural treponematosis of the African primate. Primates Med. 1969;3: 113–121.

52. Chuma IS, Roos C, Atickem A, Bohm T, Anthony Collins D, Grillová L, et al. Strain diversity of *Treponema pallidum* subsp. *pertenue* suggests rare interspecies transmission in African nonhuman primates. Sci Rep. 2019;9: 14243. doi:10.1038/s41598-019-50779-9

53. Parks DH, Porter M, Churcher S, Wang S, Blouin C, Whalley J, et al. GenGIS: A geospatial information system for genomic data. Genome Res. 2009;19: 1896–1904. doi:10.1101/GR.095612.109

54. Davies TJ, Pedersen AB. Phylogeny and geography predict pathogen community similarity in wild primates and humans. Proceedings Biol Sci. 2008;275: 1695–701. doi:10.1098/rspb.2008.0284

55. Kumm HW, Turner TB. The transmission of yaws from man to rabbits by an insect vector, *Hippelates pallipes*. Am J Trop Med Hyg. 1936;s1-16: 245–271. doi:https://doi.org/10.4269/ajtmh.1936.s1-16.245

56. Satchell GH, Harrison RA. Experimental observations on the possibility of transmission of yaws by wound-feeding Diptera, in Western Samoa. Trans R Soc Trop Med Hyg. 1953;47: 148–153.

57. Gogarten JF, Düx A, Mubemba B, Pléh K, Hoffmann C, Mielke A, et al. Tropical rainforest flies carrying pathogens form stable associations with social nonhuman primates. Mol Ecol. 2019; mec.15145. doi:10.1111/mec.15145

58. Knauf S, Raphael J, Mitjà O, Lejora IAV, Chuma IS, Batamuzi EK, et al. Isolation of Treponema DNA from necrophagous flies in a natural ecosystem. EBioMedicine. 2016;11: 85–90. doi:10.1016/j.ebiom.2016.07.033

59. Stamm L V. Flies and Yaws: Molecular studies provide new insight. EBioMedicine. 2016;11: 9–10. doi:10.1016/j.ebiom.2016.08.028

60. Boesch C, Boesch H. Hunting behavior of wild chimpanzees in the Taï National Park. Am J Phys Anthropol. 1989;78: 547–573. doi:10.1002/ajpa.1330780410

61. Gogarten JF, Akoua-Koffi C, Calvignac-Spencer S, Leendertz SAJ, Weiss S, Couacy-Hymann E, et al. The ecology of primate retroviruses – An assessment of 12 years of retroviral studies in the Taï national park area, Côte d׳Ivoire. Virology. 2014;460–461: 147–153. doi:10.1016/J.VIROL.2014.05.012

62. Wallis J& RL. Primate Conservation: The prevention of disease transmission. Int J Primatol. 1999;20: 803–826. doi:https://doi.org/10.1023/A:1020879700286

63. Fabricius T, Winther C, Ewertsen C, Kemp M, Nielsen SD. Osteitis in the dens of axis caused by *Treponema pallidum*. BMC Infect Dis. 2013;13: 347. doi:10.1186/1471-2334-13-347

